# Shape-Shifting Conotoxins Reveal Divergent Pore-Targeting Mechanisms in Nicotinic Receptors

**DOI:** 10.1101/2025.04.23.650119

**Authors:** Biddut Bhattacharjee, Colleen M. Noviello, Md Mahfuz Rahman, John P. Mayer, Joanna Gajewiak, J. Michael McIntosh, Ryan E. Hibbs, Michael H. B. Stowell

**Affiliations:** Department of Molecular, Cellular & Developmental Biology, University of Colorado, Boulder, CO, USA; Department of Neurobiology, and Department of Pharmacology, University of California San Diego, La Jolla, CA, USA; George E. Wahlen Veterans Affairs Medical Center, and Department of Psychiatry and School of Biological Sciences, University of Utah, Salt Lake City, UT, USA

**Keywords:** α-conotoxins, nicotinic acetylcholine receptor, conformational variability, targeted evolution

## Abstract

The neuronal α7 nicotinic acetylcholine receptor (α7-nAChR) and muscle-type nicotinic acetylcholine receptor (mt-nAChR) are pivotal in synaptic signaling within the brain and the neuromuscular junction respectively. Additionally, they are both targets of a wide range of drugs and toxins. Here, we utilize cryoEM to delineate structures of these nAChRs in complex with the conotoxins ImI and ImII from *Conus imperialis*. Despite nominal sequence divergence, ImI and ImII exhibit discrete binding preferences and adopt drastically different conformational states upon binding. ImI engages the orthosteric sites of the α7-nAChR, while ImII forms distinct pore-bound complexes with both the α7-nAChR and mt-nAChR. Strikingly, ImII adopts a compact globular conformation that binds as a monomer to the α7-nAChR pore and as an oblate dimer to the mt-nAChR pore. These structural characterizations advance our understanding of nAChR-ligand interactions as well as the subtle sequence variations that result in dramatically altered functional outcomes in small peptide toxins. Importantly, these results further elucidate the broad nature of cone snail toxin activities and highlight how targeted molecular evolution can give rise to functionally similar activities with surprisingly diverse mechanisms of action.

## Introduction

The venomous sea snails from the family *Conidea* produce an expansive number of peptides that target a diverse array of ion channels, G protein-coupled receptors, transporters, and enzymes as well as the insulin receptor (*1*, *2*). These conotoxins have evolved for prey capture and defense and disable a host of physiological functions.

Their remarkable potency, as well as specificity, make conotoxins a fascinating subject for neuroscientific research and are valuable starting points for drug development(*3*). Genomic and proteomic efforts have led to the identification of over 10,000 conotoxin sequences of which only a few have been thoroughly studied (*1*). Of these, the α-conotoxins have been the subject of substantial investigation due to their specificity in targeting nicotinic acetylcholine receptors (nAChRs), which are found at both the neuromuscular junction and within the central nervous system (*4*). Their exquisite subtype selectivity and small size have led to the development of analogs as potential drug candidates (*5*) including the FDA-approved pain medication ziconotide (*6*, *7*).

Additionally, from an evolutionary perspective, the toxin genes themselves display enhanced duplication and rapid evolution rates that greatly exceed other protein classes (*8*). This rapid evolution ensures that the *Conidea* outpace the prey in the biological arms race between hunter and hunted. Intriguingly, the mechanism of these enhanced rates is proposed to arise from a combination of point mutations, indels, recombination, and alternative splicing events. Much of these genetic changes are driven by the preferred use of error prone DNA polymerases that selectively replicate toxin genes (*9*).

The α7 nAChR (α7-nAChR) subtype is a homopentameric channel that is prominently expressed in the central nervous system where it plays a modulatory role (*10*, *11*) and is linked to neurological disorders (*12*) and degenerative diseases such as Alzheimer’s (*13*, *14*). Muscle-type nAChRs (mt-nAChR), typified by those found in *Torpedo californica*, are heteromeric and essential for neuromuscular junction signal transduction. They represent the archetypal Cys-loop ligand-gated ion channel and were the first identified and studied. The structures of both human α7-nAChR and *Torpedo* mt-nAChR in a resting α-bungarotoxin (α-Btx) bound state and ligand-bound desensitized state have been previously reported (*15–17*) as has the fetal and adult mt-AChR from bovine muscle (*18*). To gain molecular insight into conotoxin channel specificity that has driven selection, we investigated the α-conotoxins (α-Ctx) ImI and ImII bound to both the α7-nAChR and mt-nAChR. ImI and ImII have high amino acid sequence identity (75%) and form a distinct amino acid sequence clade of conotoxins (**fig 1A)**. Despite their high sequence identity (**fig 1B)**, and similarity in targeting nAChRs, they display markedly different pharmacology. While ImI inhibits homomeric α7-nAChRs, ImII inhibits both α7 and mt-nAChRs (*19–22*). These combined properties make them intriguing targets for structure-function studies as well as fascinating examples of the process of natural selection. Both toxins contain two disulfide bonds forming a compact rigid structure as observed for ImI and presumably for ImII, although no structure of ImII has been reported (**fig 1C)**. Here we report the cryoEM structures of ImI bound to the α7-nAChR and ImII bound to both the α7-nAChR and mt-nAChR. Remarkably, we observe that ImII displays substantial conformational variability in binding, adopting a globular ImI-like state upon binding to α7-nAChR and an oblate dimeric state upon binding to mt-nAChR. These observations present a remarkable example of extraordinary mechanistic diversity. Given the vast number of conotoxins (*1*, *3*), we expect that other intriguing examples of this mechanistic diversification through natural selection are yet to be discovered.

**Figure 1:**
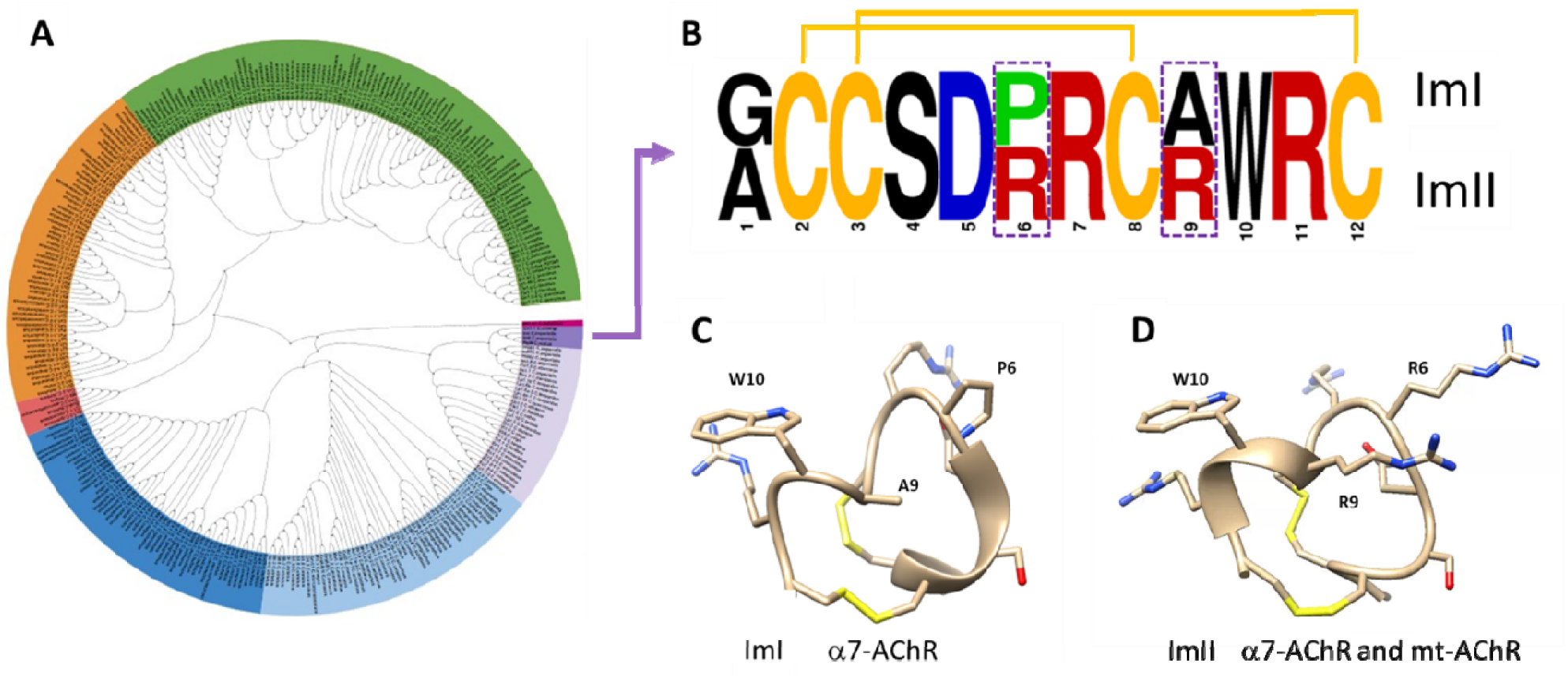
Cladogram, sequence and structures of ImI and ImII. **A)** Cladogram of the A-conotoxin family. The ImI/ImII subclade is colored in mauve, the Ch1.1 colored in light mauve has the identical mature toxin sequence as ImI but a different precursor sequence. **B)** Sequence comparison of ImI and ImII, the C-terminal cysteine (Ctx-Cys12) in both toxins is amidated. **C)** ImI structure as observed bound to AChBP (PDB: 2BYP). **D)** ImII structure as predicted by AlphaFold3. Ctx-Trp10 and the major mutations at positions 6 and 9 are labelled.

### ImI-α7-nAChR Complex

Foundational functional studies of ImI identified two regions crucial for high-affinity binding to neuronal α7 receptors (*21*). The first key region is the triad Asp-Pro-Arg in the first loop of ImI, with the Pro6 residue playing a pivotal role in maintaining structural rigidity presumed essential for the toxin’s bioactivity. Mutations of Pro6Gly, Asp5Glu, and Arg7Lys, along with charge inversions at positions 5 and 7, significantly reduce ImI’s affinity, indicating that both charge and side chain length at these positions are vital (*23*). The second crucial region is the single tryptophan at position 10 in the second loop (*23*). While mutations in the non-cysteine residues of this loop generally do not affect the affinity of ImI, replacing tryptophan with threonine markedly diminishes affinity, underscoring the importance of an aromatic side chain at this position for stabilizing the ImI-α7-nAChR receptor complex. These studies highlight the nuanced molecular interactions critical for ImI targeting the α7-nAChR. The structure of ImI bound to the acetylcholine binding protein (AChBP) provided the first structural details of this toxin interacting with the orthosteric ligand binding site (*24*), but did not provide structural insight into the state of the receptor pore. Electrophysiological studies on α7-nAChR conflict in their assessment of mechanisms of inhibition; one study suggested ImI stabilizes a desensitized state of the receptor (*24*), but others conclude it acts more as a competitive antagonist to stabilize a resting-like state(*25*).

The cryoEM structure of α7-nAChR-ImI complex determined here shows that the toxin is tightly bound to the ligand-binding domain, interfacing with key aromatic residues that are crucial for acetylcholine recognition (**fig 2A)**. While the conformation of ImI and its interactions at the binding site are conserved overall with the AChBP structure (**fig 2B**), there are several nuanced differences. In the α7-nAChR structure, the toxin forms a series of hydrogen bonds with the glucose moiety of the N-glycosylated Asn110, that further stabilizes the interaction. This was not observed for the AChBP bound structure, as AChBP lacks this key glycosylated residue. The conserved, buried Ctx-Arg7 does not form a canonical π-cation interaction with Tyr92, but rather forms a hydrogen bond between the phenolic hydroxyl and the Ctx-Arg7 epsilon guanidinium nitrogen. Additionally, Ctx-Arg11 adopts an alternative rotamer and forms an unusual stacking interaction with the loop C vicinal disulfide bond, as well as a hydrogen bond between the Tyr194 phenolic hydroxyl and the epsilon guanidinium nitrogen like that seen between Ctx-Arg7 and Tyr92. The vicinal disulfide Ctx-Arg11 interaction is consistent with a cysteine n→π* interaction (*26*), wherein the n_s_ and n_p_ orbitals of the disulfide donate electrons into the CN double bond π* orbital of Ctx-Arg11 (**fig S3A**). Importantly, and contrary to a prior study suggesting that ImI stabilizes the desensitized state (*24*), the ImI complex adopts a closed resting pore conformation nearly identical to the α-Btx bound state (**fig 2D-E**) wherein loop C is in the displaced conformation (*15*). Thus, ImI binds to the orthosteric site and stabilizes the resting state of the receptor, similar to, but not identical to the α−Btx bound state.

**Figure 2:**
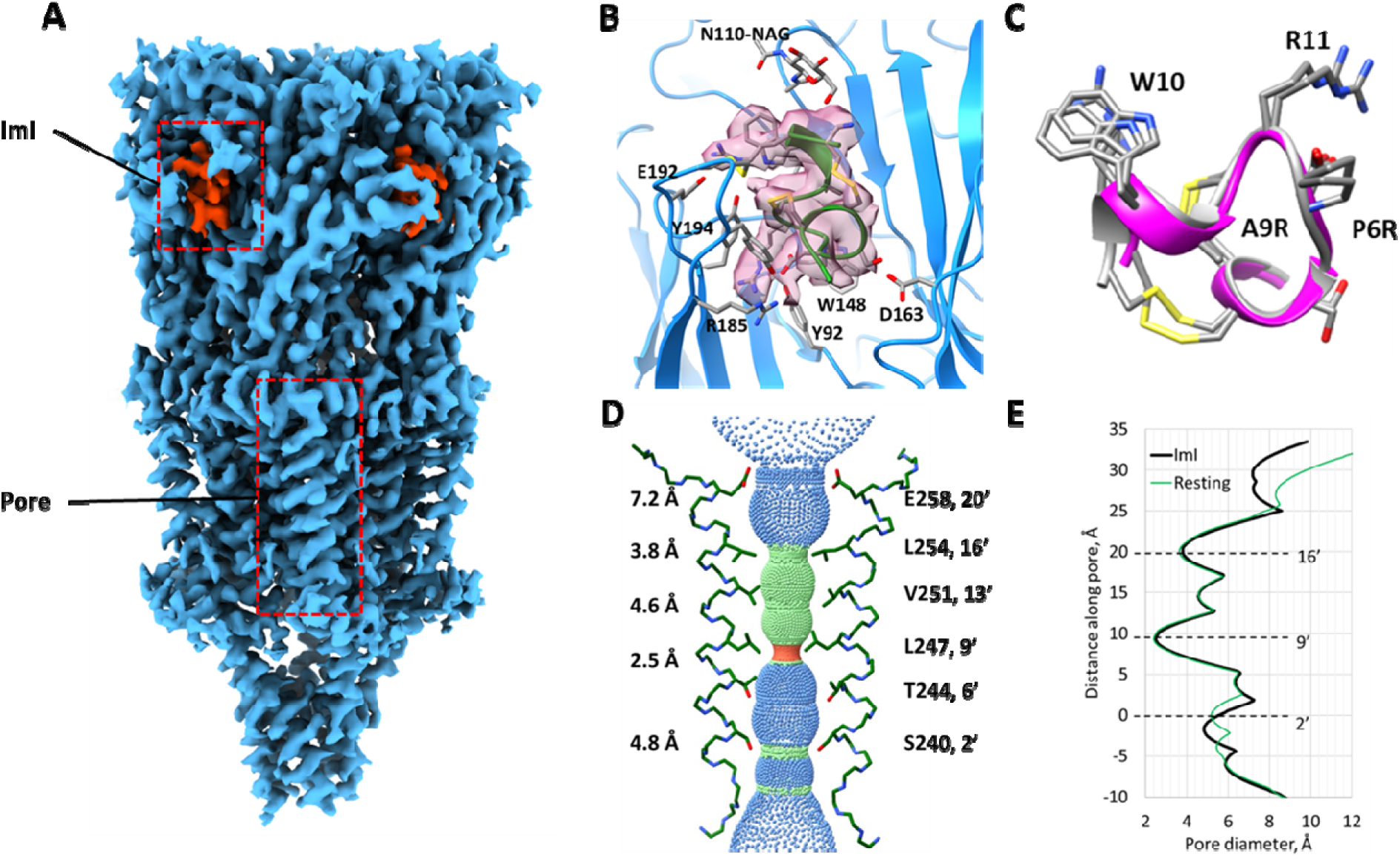
ImI stabilizes a resting closed state of human α7-nAChR. **A)** View along the membrane plane of the ImI-α7-nAChR cryoEM map, the ion pore and ImI binding site are outlined in hashed red. **B)** Molecular detail of the ImI binding pocket. The cryoEM density is shown in transparent mauve and the principle ImI contact residues are labeled. **C)** Structural comparison of ImI bound to α7-nAChR and AChBP (PDB 2C9T). **D)** Pore structure of the ImI complex structure showing the major constrictions and corresponding diameters. **E)** Pore comparison of the ImI structure to the a-bungarotoxin structure (PDB:7KOO).

Prior work combined mutational and docking studies to determine key residues responsible for selectivity of ImI over ImII, and showed that they bind at separate sites on the receptor (*19*, *20*). This work utilized homology models of α7-nAChR for the docking analysis and we performed similar calculations using Rosetta (*27*, *28*) and the cryoEM structure of the ImI bound complex, to better understand the site selectivity of ImI versus ImII. This analysis revealed that the major ImII substitutions, Pro6Arg and Ala9Arg, are not tolerated energetically (**fig S1**). While this result readily explains why ImII does not bind the orthosteric site, it does not provide insight into how or where ImII engages α7-nAChR, or how it inhibits channel function.

### ImII-α7-nAChR Complex

The binding site of ImII to the α7-nAChR has been controversial. Although prior studies agree that the ImII binding site is distinct from the ImI binding site, they disagree with regard to the overlap relative to the orthosteric site (*20*, *22*). To reconcile this prior controversy, we employed cryoEM to identify the ImII binding site. Indeed, our initial maps clearly indicated that ImII does not bind near the orthosteric sites but rather occupies a central position within the pore. The cryoEM analysis was complicated by the small mass of ImII compared to α7-nAChR, combined with the symmetry mismatch (C_1_ versus C_5_) of the complex. As a result, the initial density maps were difficult to interpret. We employed several strategies to improve the quality of the map (see methods) that allowed us to build a model of ImII in a compact globular state with good geometry and a number of chemically favorable interactions with the α7-nAChR (**fig 3**). Nonetheless, while we are confident in the location and compact globular state of ImII in the pore, there are likely several stable bound states that are energetically favorable. To explore these possibilities we again implemented a Rosetta docking protocol (*27*) with ImII and α7-nAChR. This analysis showed that the majority of stable bound states of ImII are in a conformation similar to that modeled from the cryoEM map and further support our interpretation that ImII binds α7-nAChR in several related compact globular conformations (**Fig S6**).

**Figure 3:**
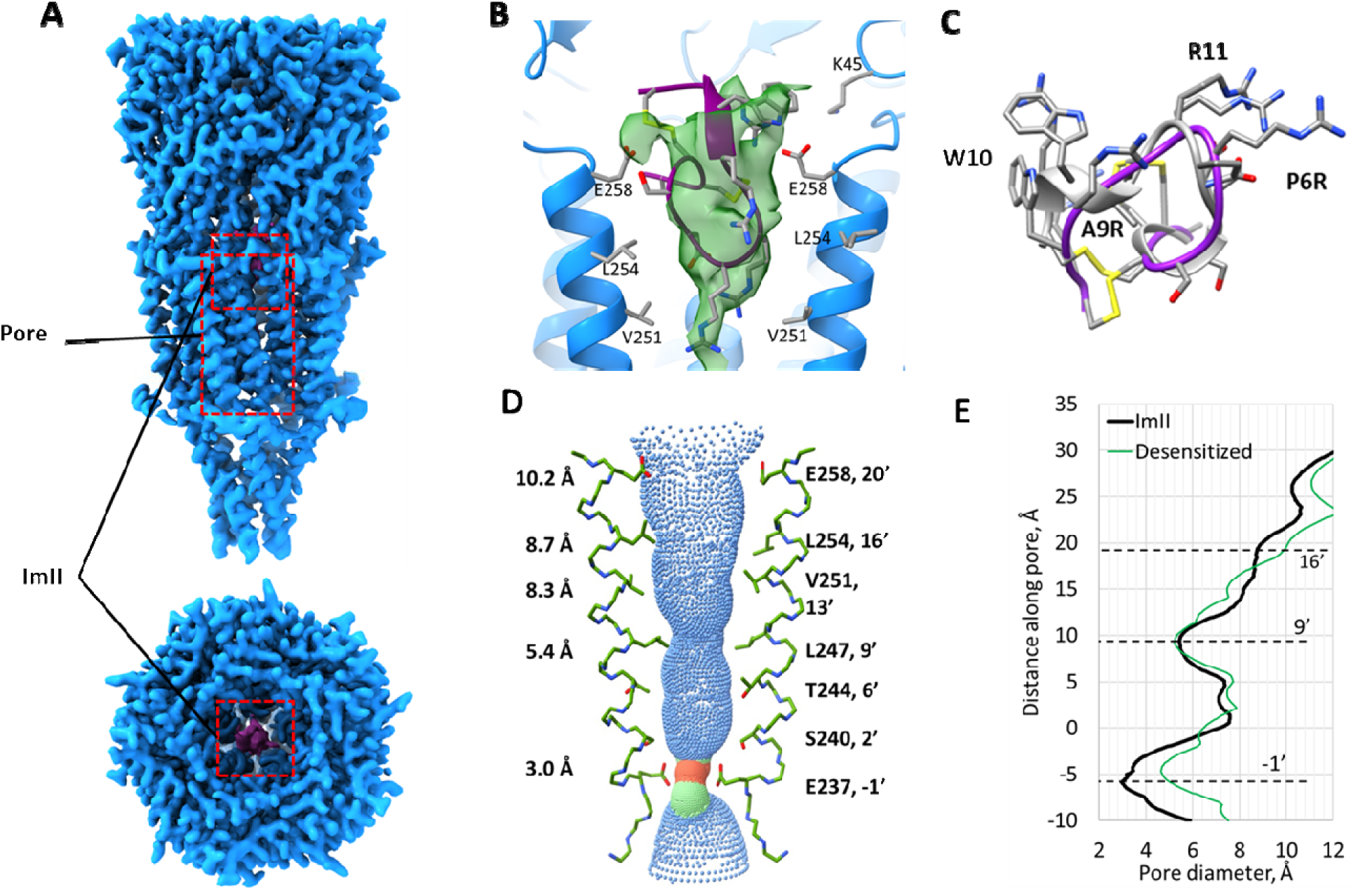
ImII binds the pore in a globular conformation and stabilizes a desensitized state of human α7-nAChR. **A)** View along the membrane plane and looking down the pore of the ImII-α7-nAChR cryoEM map, ImII is shown in red. **B)** ImII binding pocket with cryoEM density of ImII and labelled contact residues. **C)** Structural comparison of ImI and ImII bound to α7-nAChR. **D)** Pore structure of the ImII complex showing the major constrictions and corresponding diameters. **E)** Pore comparison of the ImII structure to the desensitized α7-nAChR structure (PDB:7KOQ).

Dramatically contrasting with the ImI α7-nAChR complex, the α7-nAChR receptor bound by ImII reveals the toxin lodged within the central pore, adopting a globular conformation similar but not identical to ImI bound to the orthostatic site, (**fig 3B**). The pore adopts a desensitized state (**fig 3C**), implying that the toxin binds and stabilizes an agonist-bound state as our preparation contained the high affinity agonist epibatidine.

Indeed, when we tried to obtain an ImII-bound structure in the absence of agonist, we found no occupancy of this site by ImII (data not shown). Importantly, this pore density is not seen in the epibatidine-only map previously published(*15*). The observation that ImII stabilizes a desensitized states reconciles the apparently contradicting data suggesting that ImII can compete with α−Btx in α7-nAChR binding (*22*), as ImII binding would drive the receptor into a desensitized state that has lower affinity for α−Btx binding (*29*). As a result of the subunit interdependency of α−Btx binding (*30*) a ligand such as ImII that drives α7-nAChR towards desensitization would pharmacologically appear to act in a competitive manner despite a binding site that does not physically overlap, thus ImII binding acts as an allosteric antagonist to α−Btx binding.

ImII binds in the ion-conducting pore of the α7-nAChR near the extracellular end of the transmembrane domain, interacting with the pore-lining M2 helices. Unlike ImI, where the triad Asp5-Pro6-Arg7 in the first loop of the toxin plays the dominant role in binding to the orthosteric site, the second loop of ImII contributes predominantly to the binding (**fig 3B)**. Ctx-Arg11 guanidinium forms hydrogen bonds and a salt-bridge with the sidechain carboxyls of Glu258. The indole NH moiety of Ctx-Trp10 makes a hydrogen bond with the carboxyl oxygen of Glu258. Additionally, these Ctx-Trp10 and Ctx-Arg11 are stabilized by a cation-π interaction with a centroidal distance of 4.8 Å. The positively charged amino group of Lys45 from β1-β2 loop lies at approximately 8.5 Å from the Tx-Trp10 indole, suggesting a possible cation-π interaction. The backbone nitrogens of Tx-Ala1 and Ctx-Cys2 make hydrogen bonds with the carboxyl group of Glu258. The sidechain of Ctx-Arg6 extends deeply into the ion-conducting pore toward the intracellular side of the α7-nAChR. Ctx-Asp5 and Ctx-Arg7 in the first loop of ImII form a salt-bridge between their guanidinium and the carboxyl groups while the side-chains of both Ctx-Arg6 and Ctx-Arg7 make hydrophobic contacts with Val251 and Leu254, respectively (**fig S3B)**.

### ImII-mt-nAChR Complex

The binding of ImII to the mt-nAChR was previously proposed to have a distinct binding site from the classical orthosteric site (*22*). The cryoEM structure of ImII bound to mt-nAChR shows an ImII non-covalent dimer occluding the pore and the channel in a desensitized state (**fig 4**). We term the toxin closest to the channel pore the lower ImII and the toxin furthest from the pore the upper ImII. Each monomer of the ImII dimer adopts an oblate conformation distinct from both ImI and ImII bound to the α7-nAChR (**fig 4**). While the quality of the map was sufficient to confirm that the bound ImII was in fact the globular form (2-8:3-12 linkage), due to the unusual and unexpected conformation of ImII bound to mt-nAChR we prepared all three possible isomers (globular, bead and ribbon) to further validate that the globular form of the peptide was utilized for structural analysis (see methods). The ImII dimer interaction consists primarily of several hydrogen bond interactions, van der Walls interactions, and an unusual disulfide bond-disulfide bond interaction. The intermolecular hydrogen bond interactions are formed between the backbone oxygen of Cys2 and the gamma oxygen of Ser4 from the upper ImII with the delta oxygen Asp5 from the lower ImII, as well as between the Ala1 N-terminal nitrogen of the upper ImII and the delta oxygen of Asp5 from the lower ImII. The buried surface area between the two ImII monomers is only 237 Å^2^, making this interaction unlikely to exist in solution. The disulfide bond-disulfide bond interaction is unusual and consists of the upper ImII Cys12 interacting with the lower ImII Cys3 (**fig 4B**).

**Figure 4:**
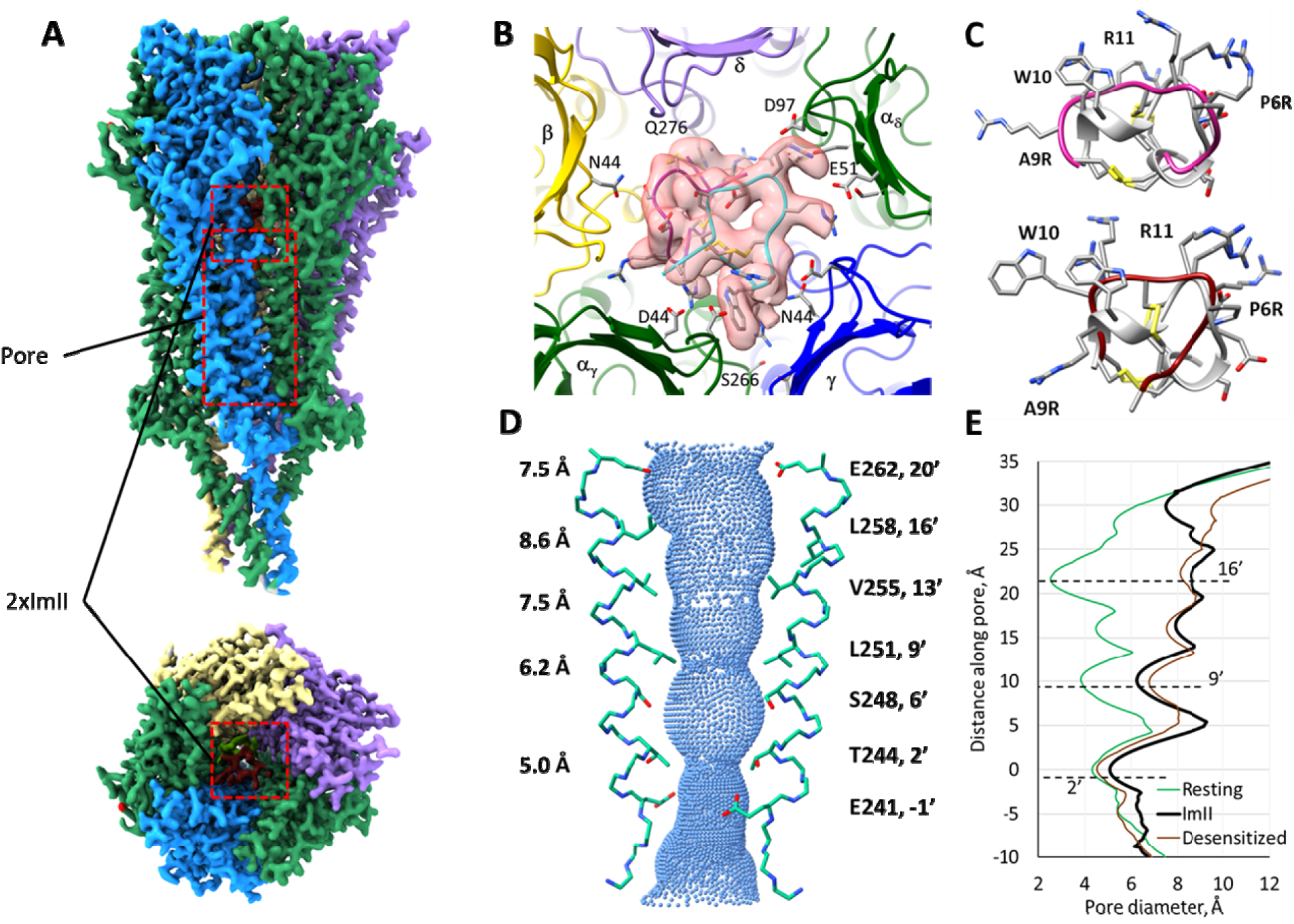
ImII binds the pore of mt-nAChR in an oblate conformation as a dimer and stabilizes a desensitized state. **A)** View along the membrane plane and looking down the pore of the ImII-mt-nAChR cryoEM map, ImII is shown in red. **B)** ImII binding pocket with cryoEM density of ImII and contact residues. **C)** Structural comparison of ImII bound to mt-nAChR and ImI (PDB 2C9T), the upper and lower bound positions are shown. **D)** Pore structure of the ImII complex showing the major constrictions and corresponding diameters. **E)** Pore comparison of the ImII structure to the resting and desensitized state structures of mt-nAChR (PDB:7SMS and 7SMM).

A high surface and electrostatic complementarity are observed between both ImII molecules and the mt-nAChR. The upper ImII has a buried surface area of 869 Å^2^ and the lower ImII has a buried surface area of 750 Å^2^. About 25% of upper ImII buried area is with the lower ImII, and the remaining 75% is with subunits a_y_ya_δ_. The upper ImII predominantly contacts β1, β2, β1-β2 loop and β3-β4 loop, and forms multiple hydrogen bonds and salt-bridges through the sidechains Ctx-Arg6, Ctx-Arg7 and Ctx-Arg11, along with other residues in the electrostatically favorable binding grooves.

The guanidinium amine of upper Ctx-Arg6 interacts with the carboxyl groups of Glu51, Asp44 and Asp97 of the a_δ_ subunit. While Ctx-Arg7 enters the narrow lateral tunnel defined by the β1 and loop F (β8-β9) of a_δ_ and the β1-β2 loop and the adjacent vertical portion of the Cys-loop of y subunit interacting with Ser42 and Glu51 of a_δ_ subunit and Glu47 of y subunit. Strikingly, the equivalent tunnel penetrated by Ctx-Arg11 and formed at the interface between a_y_ and y is closed, though much wider near the ECD vestibule, due to the M2-M3 loop from a_y_ and the F-loop from y. The Ctx-Arg11 rotamer projects deep into the tunnel forming two hydrogen bonds between its guanidinium group and a_y_ Ser266 (M2-M3 linker) and yAsn44 (β1), and one polar contact with γGlu183 in F-loop. a_y_ Asp97 carboxyl group in the β3-β4 loop stabilizes both Ctx-Arg9 and Ctx-Trp10 by interacting with nitrogen atoms where the indole sidechain is tucked into a pocket underneath the loop (**fig S4A**).

The lower ImII forms a total of 750 Å^2^ (44%) interface with subunits a_δ_ of3a_y_ while inserting residues Ctx-Arg9, Ctx-Trp10 and Ctx-Arg11 into the transmembrane pore of mt-nAChR. The partial insertion of the toxin into the pore is likely due to the sharper curvature of the second loop resulting from the hydrogen bond between the backbone carbonyl oxygen of Ctx-Cys8 and the backbone nitrogen of Ctx-Arg11, relative to the loop in the upper ImII. The Ctx-Arg11 guanidinium forms a polar bond with the negatively charged sidechain of a_δ_ Asp262 in the pore lining M2 helix. The backbone oxygen of Ctx-Trp10 engages in a hydrogen bond with the amine of δ-Gln276 of M2. On the diametrically opposite side of the ECD vestibule, Ctx-Arg6 fits into the radial tunnel formed at the interface between subunits a_y_ and β, the base of which is supported by the M2-M3 linker from β subunit. The backbone carbonyl of Ile41 in β1 and the sidechain of Glu175 in F-loop, both from a_y_, engages with the positively charged sidechain of Ctx-R6. Additional binding interactions with the lower ImII are provided by a_y_ Asp44 with Ctx-Arg7 and the βAsn44 with Ctx-Cys3 (**Fig S4B**).

## Discussion

Here we have elucidated the structural details of how two closely related conotoxins, ImI and ImII, antagonize the α7- and mt-nAChRs. Conotoxins comprise one of the largest known sets of diverse peptide toxins that target multiple physiological functions, and they offer the potential for discovering unique pharmacological compounds. The rapid evolution of toxin genes provides a fascinating example of directed evolution within a genome. Specifically, the structures reveal how minor changes in toxin sequence can lead to dramatic differences in receptor interaction and mechanism of nAChR inhibition. The profound conformational flexibility of ImII and its distinctive engagement with α7- and mt-nAChRs emphasize the dynamic nature of ligand-receptor interactions and suggest that such flexibility was selected to achieve a polypharmacological action. In addition, the non-covalent dimerization of ImII within the muscle-type nAChR pore raises interesting possibilities regarding the regulation of ion channels by bivalent or multimeric ligands and represents a new and unique conotoxin inhibition mechanism not previously observed. An intriguing question arises as to how such toxins could arise through natural selection given the unique mechanisms of action and distinct molecular target sites observed.

## Materials and Methods

### Toxin synthesis

This procedure closely follows previously published methods (*31*). All standard solvents and reagents were obtained from Sigma-Aldrich unless otherwise specified, Fmoc-protected amino acids were purchased from Vivitide, H-Rink-ChemMatrix resin (loading level: 0.47 mmoml/g) from Gyros Technologies, trifluoroacetic acid (TFA) from EMD Millipore, piperidine from Alfa-Aesar, 1-hydroxy-6-chloro-benzotriazole (6-Cl-HOBt) from Creosalus-Advanced Chemtech, tri-isopropylsilane (TIS) from TCI. Analytical chromatography was performed on an Agilent 1100 model system equipped with an auto sampler using Phenomenex columns as specified below. Preparative chromatography was conducted on a Waters Model 2525 binary pump system equipped with a Model 2487 absorbance detector. Mass spectral data was collected on a Waters Synapt G2 HDMS. ImII was assembled on 0.1 mmol scale by automated solid-phase methodology using an Applied Biosystems 431A synthesizer on H-Rink-ChemMatrix support or an AAPPTec Apex 396 synthesizer on Rink Amide MBHA resin. The standard α-fluorenyl-methoxycarbonyl/t-butyl (Fmoc/tBu) based protecting group scheme. The automated cycles utilized 6-Cl-hydroxybenztriazole (6-Cl-HOBt)/ diisopropylcarbodiimide (DIC) for activation and 20% piperidine/DMF for deprotection. N-terminal acetylation was performed manually by treating the peptide resin with a DMF solution containing 5% acetic anhydride for 30 minutes. Resin cleavage and side-chain deprotection were carried in 15 ml of TFA containing 2% v/v of the following scavengers: water, TIS, thioanisole, β-mercaptoethanol for 2.0 hours. The crude peptide was recovered by addition of excess cold diethyl ether, followed by several washes of the precipitate with diethyl ether and air drying. The three theoretical disulfide isomers were synthesized by directed disulfide bond formation using the acetamidomethyl (Acm) group to protect two of the four cysteines with the other two protected with trityl groups. Minimization of aspartimide formation for ImII was achieved using Asp(OBno) instead of Asp(OtBu) in position five. Following TFA or Reagent K cleavage and deprotection the first disulfide was formed by oxidation in 50 mM ammonium bicarbonate buffer, stirring overnight in an open beaker at 1 mg/ml or in 20 mm potassium ferricyanide and 0.1 m Tris-HCl, pH 7.5. Subsequently the Acm-protected cysteine pair was then converted to the disulfide by treatment with iodine and acetic acid and the product purified by preparative HPLC to >95% and mass confirmed by HRMS. The Acm-protecting scheme was necessary as the random fold of the unprotected peptide produced the “ribbon” isomer with the 2-12:3-8 linkage, and not the desired “globular” 2-8:3-12 isomer. This was confirmed by performing co-injections of the random fold with all three isomers produced using the Acm protection method, **Figure S2**. This unequivocally established the randomly folded material as having the 2-12:3-8 disulfide linkage or ribbon form.

### H. sapiens α7-nAChR expression and purification

The human α7 receptor was expressed and purified as done previously (*15*). Briefly, after transduction with Bacmam virus, HEK293S cells were cultured at 37°C for 72h before harvesting by centrifugation. Cells were lysed by passage through an Avestin Emulsiflex at 5,000-15,000 psi. Lysed cells were first centrifuged at low speed (10,000 g) to remove nuclei and unlysed cells, then centrifuged at 186,000 g for 2 h to pellet membranes. Membrane pellets were stored at −80°C until used. For purification, approximately 5g of membranes were Dounce homogenized in TBS + 1 mM PMSF and n-dodecyl-β-D-maltoside (DDM; Anatrace) was added to a final concentration of 40 mM, and solubilization was performed at 4°C while nutating. Insoluble material was then removed by centrifugation at 186,000g for 40 min. The supernatant containing solubilized protein was bound to Strep-Tactin Superflow high-capacity resin (IBA Life Sciences) via gravity flow. The resin was washed with TBS plus 1 mM DDM, supplemented with 10 µM Soy Polar Lipids (Avanti) and 0.25% w/vol cholesterol (Sigma) to promote receptor stability. The α7 receptor protein was eluted in the same wash buffer supplemented with 5 mM desthiobiotin (Sigma).

The α7-nAChR protein was concentrated to ∼500 μl, and then saposin (*32*) and soy lipids/25% cholesterol (w/w) were added to the purified protein at a molar ratio of 1:50:230 of α7-nAChR:saposin:lipid. This mixture was nutated at 4°C for 30 min before the removal of detergent by the addition of bio-beads and continued overnight at 4°C, after which incorporation was assayed by fluorescence-detection size-exclusion chromatography(*33*). The receptor-lipidic nanodisc mixture was further polished by size exclusion chromatography using only TBS as a mobile phase. For the ImI:α7 structure, the receptor was purified in the absence of ligand and ImI at 100 µm was added after concentration to ∼4-6 mg/ prior to subsequent plunge freezing for cryoEM. For the ImII:α7 structure, an initial attempt with the Apo preparation failed to yield any noticeable density for ImII. Thus, the agonist epibatidine (TOCRIS) was included at 20 µm throughout the prep along with 4 mM EGTA to bias the receptor towards a desensitized state. Then 100 µm ImII was added after SEC to pooled, concentrated fractions prior to freezing grids.

### T. californica mt-nAChR purification

Nicotinic acetylcholine receptor protein was purified from the electric organs of *Tetronarce californica* (EastCoast Bio) as previously described (*16*, *17*, *34*). Briefly, 100 g of frozen tissue were thawed in 300 mL of buffer A (20 mM NaH_2_PO_4_, 400 mM NaCl, pH 7.4) supplemented with 150 mg NEM (N-ethylmaleimide, sigma). Thawed tissue was homogenized and centrifuged at 3220 g for 15 min. The supernatant was passed through cheesecloth, and a protease inhibitor tablet (Complete mini, Sigma) was added to it with gentle stirring. Cell membranes were pelleted by centrifuging this supernatant at 105,000 g for 30 min. The membrane pellet was collected and resuspended in buffer B (20 mM Tris, 80 mM NaCl, 1 mM EDTA, 20% sucrose, pH 11.0) and incubated at 4 °C for 30 min. The resuspension was centrifuged at 105,000 g for 30 min. The membrane pellet was washed twice with buffer C (20 mM NaH_2_PO_4,_ 80 mM NaCl, pH 7.4) and stored at −80°C until further use. For protein purification, 2 g of the membrane pellet were thawed on ice. A Dounce homogenizer was used to resuspend the pellet in 50 mL of buffer C. Triton X-100 (1.5% v/v) and phenylmethylsulfonyl fluoride (PMSF, 1 mM) were added to the sample and gently rocked for an hour in at 4 °C. The solubilized supernatant was collected by centrifuging the sample at 105,000 g for 30 min at 4 °C. 100 mL of buffer C were added to the supernatant. The ATM (2-[(4-aminobutanoyl)amino]-*N*,*N*,*N*-trimethylethanaminium) affinity ligand was synthesized and coupled to NHS activated Sepharose resin (Cytiva) as reported (*34*). Affinity resin (5 mL packed bed) was equilibrated with buffer B and mixed with diluted supernatant. The suspension was nutated for 1 h at 4 °C. Unbound sample was removed by washing the resin with buffer D (20 mM Tris, 80 mM NaCl, 1 mM EDTA, 1 mM *n*-dodecyl-β-D-maltoside, DDM, Anatrace, pH 7.4) and protein elution was accomplished with 50mM carbochol.

The mt-nAChR protein was concentrated to ∼4-6 mg/mL and mixed with soy polar lipids (Avanti Polar) and saposin (*32*) at a molar ratio of receptor:saposin:lipid of 1:25:150 and incubated for ∼ 5 minutes. Approximately 120 mg of Bio-Beads (SM2, Bio-Rad) were washed with methanol once, three times with Milli-Q water and once with Tris-buffered saline (80 mM NaCl, 20 mM Tris pH 7.4). Then 500–600 µl of the reaction sample above was mixed with the washed Bio-Beads and nutated overnight at 4°C. A fresh aliquot of washed Bio-Beads was added in the morning and the sample nutated for an additional hour. The saposin-reconstituted receptors were separated from the Bio-Beads using a syringe equipped with a 32-gauge needle and the passthrough sample was centrifugation at 98,600g for 40 min to remove any precipitant. The samples were then passed through a SEC column (Superose 6 10/300 GL Increase) equilibrated in Tris-buffered saline to remove empty nanodiscs from the sample. The best fractions were concentrated to ∼7 mg/mL before the addition of ImII at a concentration of 100 µM prior to plunge freezing for cryoEM.

### CryoEM sample preparation

For preparation of cryoEM grids, 200-mesh Cu 1.2/1.3 grids from Quantifoil were glow-discharged and supplemented with 0.5 mM fluorinated fos-choline-8 (Anatrace) immediately before freezing to induce random particle orientations. Typically, 3-4 μl of protein sample was applied to grids before blotting for 3.5 s at 100% humidity and 4°C. Grids were plunge frozen into liquid ethane with a Mark IV Vitrobot (FEI/ThermoFisher). Frozen grids were stored in liquid nitrogen until use for data collection.

### CryoEM data collection and processing

Samples were screened on a 200kV Talos Arctica at UT Southwestern Medical Center or a 200KV F20 at the Boulder EM Services Center and final datasets were collected on a 300 kV Titan Krios G2 (FEI) at the Pacific Northwest Center for Cryo-EM (PNCC) or a 300kV Titan Krios G1 (FEI) at the Purdue CryoEM Center, **Table 1**. The data were processed using CryoSPARC (*35*) v.4.5 using built in patch motion correction and CTF determination. Images with a CTF resolution fit worse than 8Å were discarded. Initial particles were picked using template picking and bad particles were removed by several rounds of 2D classification. Particles from 2D classes with high-resolution features were used to generate a de novo initial model for 3D classification. The 3D class with visible ICD density was low-pass filtered and used as an initial model for 3D refinement followed by several rounds of heterogeneous refinement and subsequent non-uniform refinement. Template-based motion correction was performed on the final particle data set and a final non-uniform refinement was performed with appropriate imposed symmetry, C1 for mt-nAChR, C5 and ImI-α7-nAChR. In the case of the ImII-α7-nAChR refinement, 3D classification with C1 symmetry was performed with a focused mask encompassing the toxin binding site. The resultant maps were manually inspected and rotated by increments of 72° to maximize the overlap of the asymmetric toxin density. Particle sets following rotation were combined and non-uniform 3D refinement was performed with relaxed C5 symmetry. To enhance the quality of the ImII density, local refinement was performed with a mask encompassing the toxin density and the TM domain, (**fig S7**). maps were locally sharpened using the calculated local resolution and used for subsequent model building and refinement. The final resolution, particle orientations and local resolution of the maps are shown in **fig S5**.

**Table 1.**
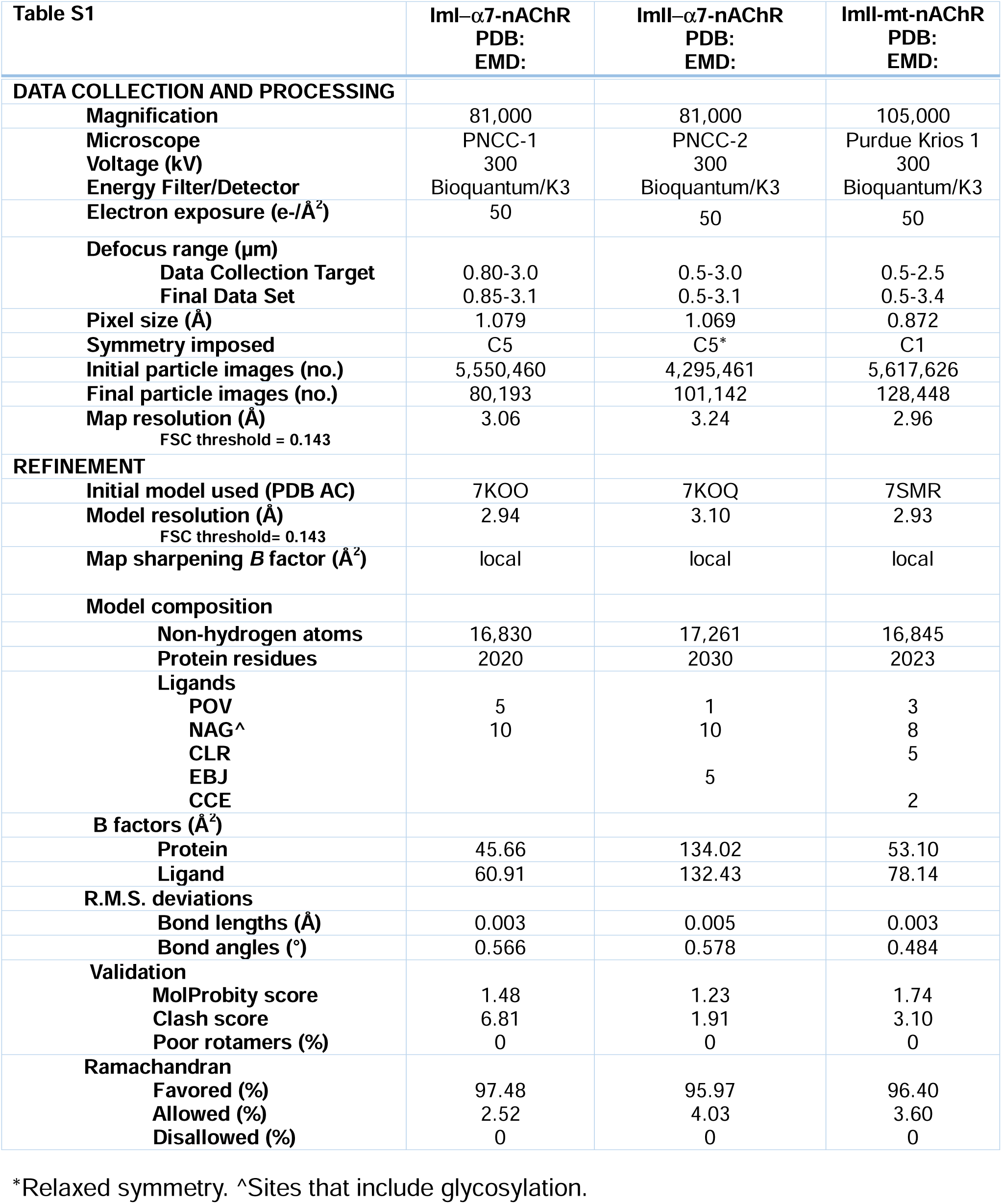
CryoEM data collection, refinement, and validation statistics. *Relaxed symmetry. ^Sites that include glycosylation.

| Table S1 | ImI-α7-nAChR<br>PDB:<br>EMD: | ImII-α7-nAChR<br>PDB:<br>EMD: | ImII-mt-nAChR<br>PDB:<br>EMD: |
| --- | --- | --- | --- |
| **DATA COLLECTION AND PROCESSING** |  |  |  |
| **Magnification** | 81,000 | 81,000 | 105,000 |
| **Microscope** | PNCC-1 | PNCC-2 | Purdue Krios 1 |
| **Voltage (kV)** | 300 | 300 | 300 |
| **Energy Filter/Detector** | Bioquantum/K3 | Bioquantum/K3 | Bioquantum/K3 |
| **Electron exposure (e-/Å<sup>2</sup>)** | 50 | 50 | 50 |
| **Defocus range (µm)** |  |  |  |
| **Data Collection Target** | 0.80-3.0 | 0.5-3.0 | 0.5-2.5 |
| **Final Data Set** | 0.85-3.1 | 0.5-3.1 | 0.5-3.4 |
| **Pixel size (Å)** | 1.079 | 1.069 | 0.872 |
| **Symmetry imposed** | C5 | C5* | C1 |
| **Initial particle images (no.)** | 5,550,460 | 4,295,461 | 5,617,626 |
| **Final particle images (no.)** | 80,193 | 101,142 | 128,448 |
| **Map resolution (Å)**<br>FSC threshold = 0.143 | 3.06 | 3.24 | 2.96 |
| **REFINEMENT** |  |  |  |
| **Initial model used (PDB AC)** | 7KOO | 7KOQ | 7SMR |
| **Model resolution (Å)**<br>FSC threshold= 0.143 | 2.94 | 3.10 | 2.93 |
| **Map sharpening B factor (Å<sup>2</sup>)** | local | local | local |
| **Model composition** |  |  |  |
| **Non-hydrogen atoms** | 16,830 | 17,261 | 16,845 |
| **Protein residues** | 2020 | 2030 | 2023 |
| **Ligands** |  |  |  |
| POV | 5 | 1 | 3 |
| NAG^ | 10 | 10 | 8 |
| CLR |  |  | 5 |
| EBJ |  | 5 |  |
| CCE |  |  | 2 |
| **B factors (Å<sup>2</sup>)** |  |  |  |
| Protein | 45.66 | 134.02 | 53.10 |
| Ligand | 60.91 | 132.43 | 78.14 |
| **R.M.S. deviations** |  |  |  |
| **Bond lengths (Å)** | 0.003 | 0.005 | 0.003 |
| **Bond angles (°)** | 0.566 | 0.578 | 0.484 |
| **Validation** |  |  |  |
| **MolProbity score** | 1.48 | 1.23 | 1.74 |
| **Clash score** | 6.81 | 1.91 | 3.10 |
| **Poor rotamers (%)** | 0 | 0 | 0 |
| **Ramachandran** |  |  |  |
| **Favored (%)** | 97.48 | 95.97 | 96.40 |
| **Allowed (%)** | 2.52 | 4.03 | 3.60 |
| **Disallowed (%)** | 0 | 0 | 0 |

### Model building, refinement and validation

The α-Bungarotoxin bound structure of the α7-nAChR (PDB 7KOO) was used as a template for initial modeling of the ImI-α7-nAChR structure, the epibatidine bound desensitized state structure of α7-nAChR (PDB 7KOQ) was used as template for initial modelling of the ImII-α7-nAChR structure and the carbochol bound desensitized state of the mt-nAChR (PDB 7SMR) was used for initial modelling of the ImII-mt-nAChR structure. The initial structure of ImI was adopted from PDB 2C9T and ImII was built from the ImI model in Chimera. Iterative cycles of manual building in COOT (*36*) with global real space refinement in Phenix (*37*) resulted in excellent stereochemistry as assessed by Molprobity (*38*). Model-map correlation was assessed using Phenix validation tools. Pore diameters were calculated using HOLE (*39*) and ligand-binding sites were analyzed by LigPlot+ (*40*). Map and structural figures were generated using UCSF Chimera (*41*) and UCSF ChimeraX (*42*).

### Rosetta Tox Dock analysis

The refined ImI-α7-nAChR structural model was used as the starting point for the substitution tolerance test. All ImI models were removed except one for the analysis. The analysis was performed using the Tox Dock server(*28*) following the procedures previously described(*27*). Similarly, the refined ImII-α7-nAChR structural model was used as the starting point for the ImII-α7-nAChR analysis.

### Data Availability

All CryoEM maps and atomic model coordinates have been deposited in the EMDB and RCSB, respectively, and will be released upon publication. PDB and EMDB accession numbers are: 9NX2/EMD-49899 mt-nAChR-ImII, 9NX1/EMD-49898 alpha7-nAChR-ImII, and 9NX0/EMD-49897, alpha7-nAChR-ImI.

All software packages used in this study are publicly available **Supplementary Information** is available in the online version of the paper.

## Acknowledgments

A portion of this research was supported by NIH grant U24GM129547 and performed at the PNCC at OHSU, U24GM116789 and performed at the Purdue CryoEM Facility. We thank Harry Scott and Thomas Klose for their assistance in cryoEM data collection. This work was supported by grants from the NIH (R01NS120496 to R.E and M.H.B.S, R35 GM136430 to J.M.M.) and the MCDB Neurodegenerative Disease Fund to M.H.B.S.

## Author Contributions

MR and CN performed the sample preparation. BB, CN, MR and MHBS performed data processing for cryoEM, structural analysis, model building and validation. JMM, JG and JPM performed peptide synthesis and analytical analysis. BB drafted the manuscript with REH and MHBS. The manuscript was revised with input from all authors.

---

**Figure S1:**
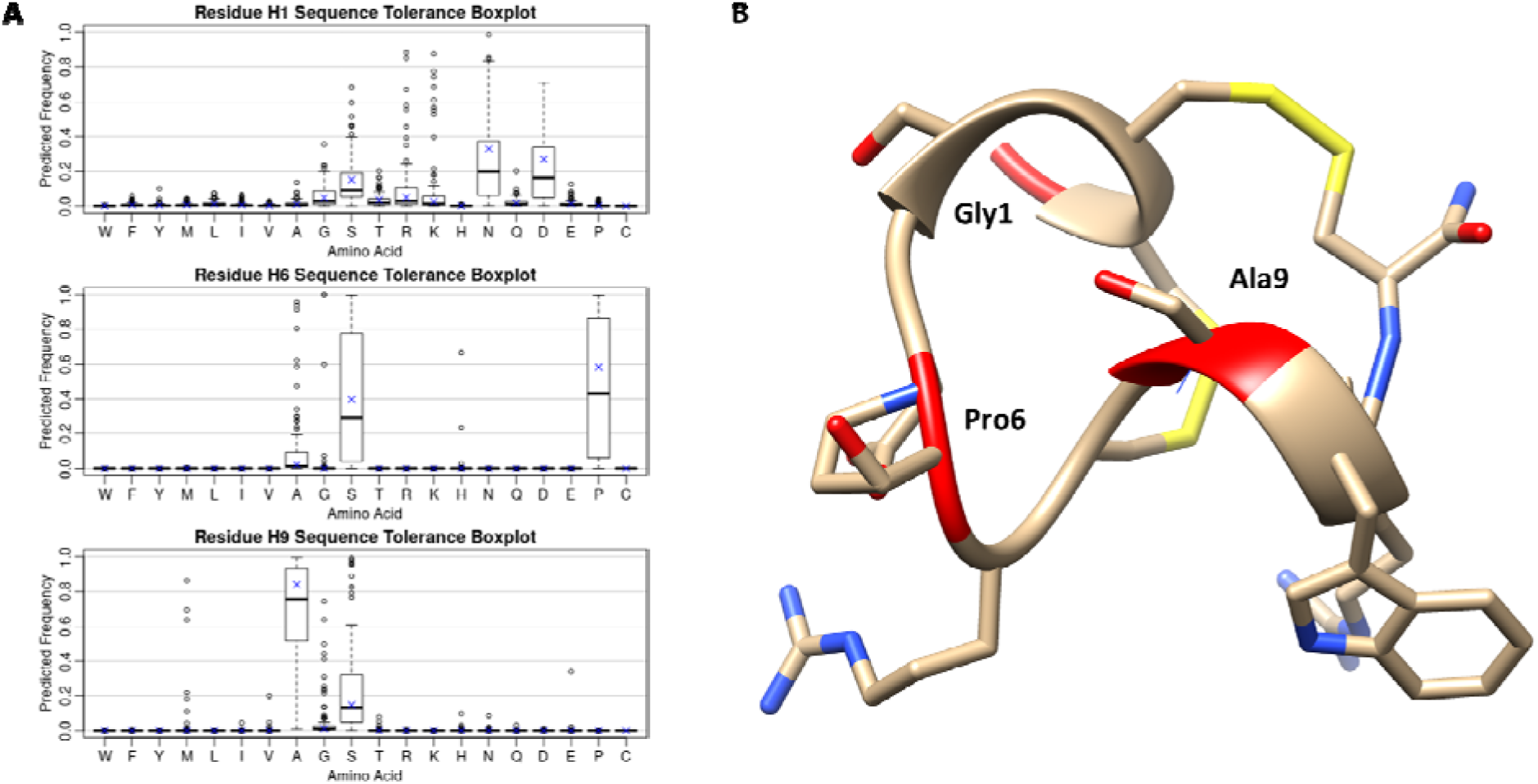
Tox Dock substitution profiling of the ImI-α7-nAChR. **A)** Tox Dock predicted frequency for substitutions in ImI bound to α7-nAChR. **B)** Structure of ImI with positions analyzed for substitution in red, Gly1, Pro6 and Ala9. The most tolerated substitution is shown overlayed with the ImI residue, i.e. serine at position 6 and 9.

**Figure S2:**
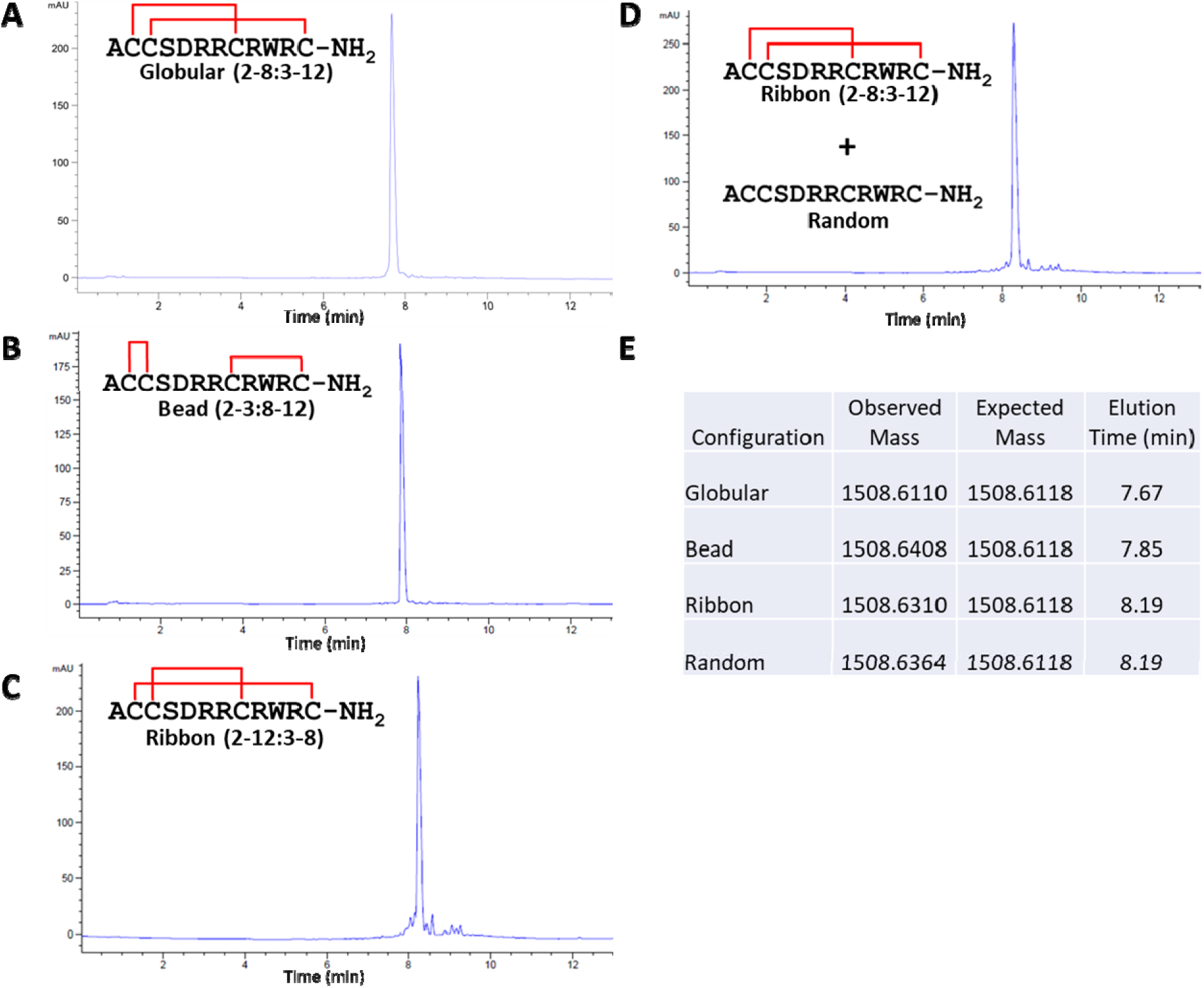
Synthesis of the 3 possible isomers of ImII and confirmation via HPLC co-injection that the random fold produces the globular isomer. **A)** HPLC trace of the globular form of ImII. **B)** HPLC trace of the bead form of ImII. **C)** HPLC trace of ribbon form of ImII. **D)** Co-injection of the ribbon form and the random fold of ImII. **E)** HR-MS and elution times of the various forms of ImII.

**Figure S3:**
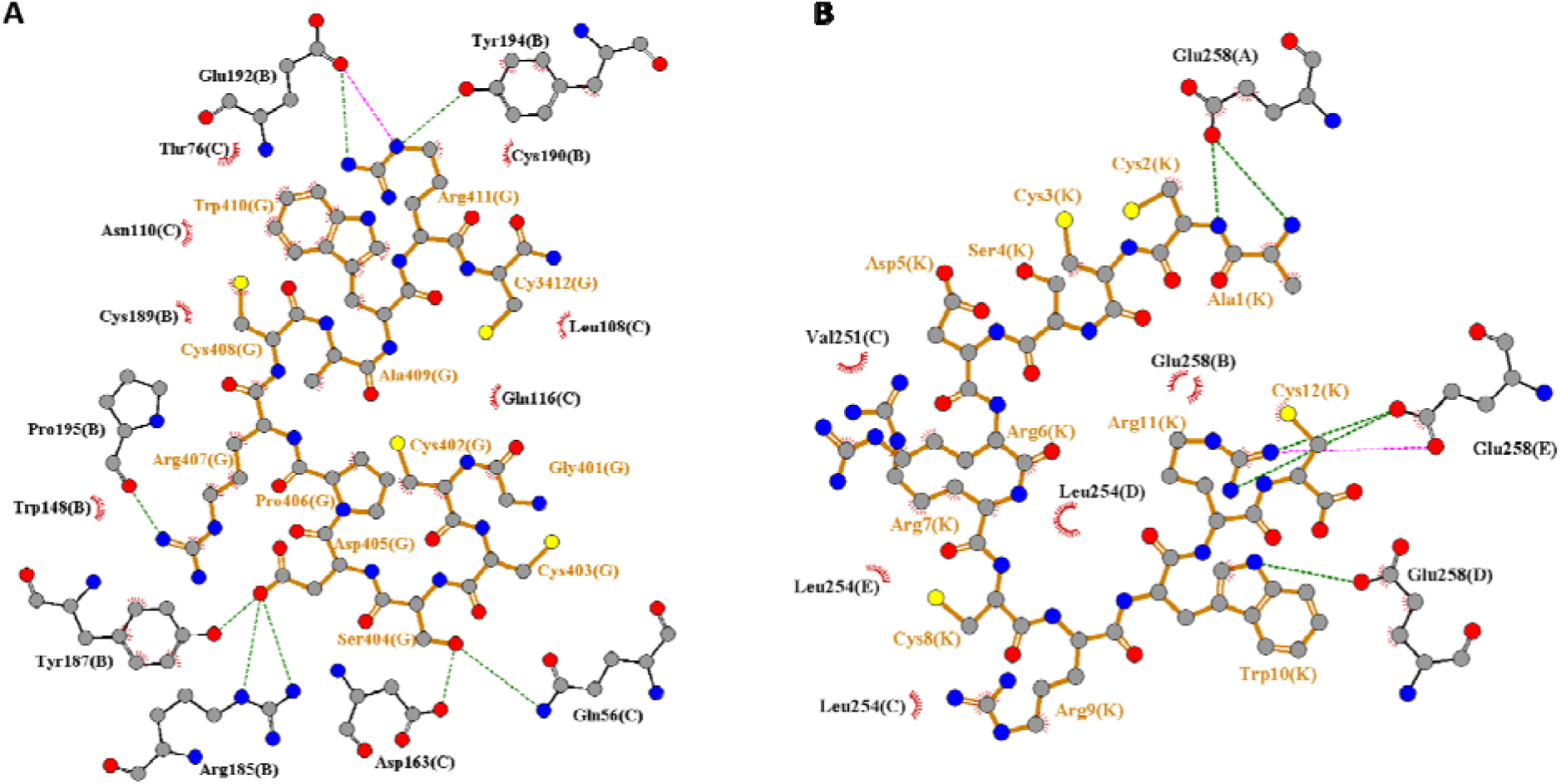
ImI and ImII interactions with α7-nAChR. **A**) ImI interaction with α7-nAChR. **B**) ImII interaction with α7-nAChR. Color coding: green denotes hydrogen bond, pink denotes salt-bridge, eyelash denotes hydrophobic interaction. The parenthetical letters correspond to the PDB chain designation.

**Figure S4:**
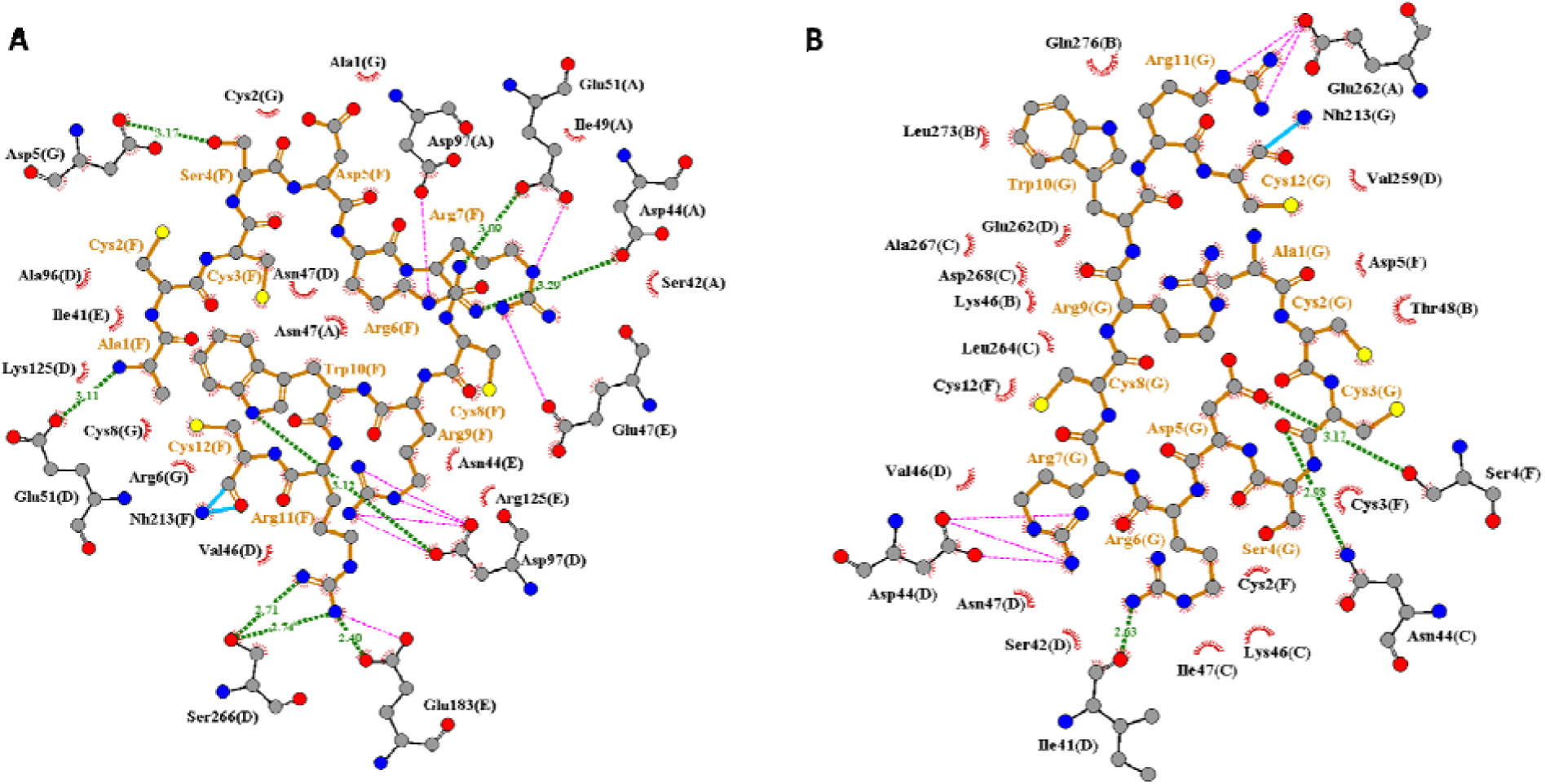
ImII interactions with mt-nAChR. **A**) Upper ImII interactions with mt-nAChR. **B**) Lower ImII interactions with mt-nAChR. Color coding: green denotes hydrogen bond, pink denotes salt-bridge, eyelash denotes hydrophobic interaction. The parenthetical letters correspond to the PDB chain designation.

**Figure S5:**
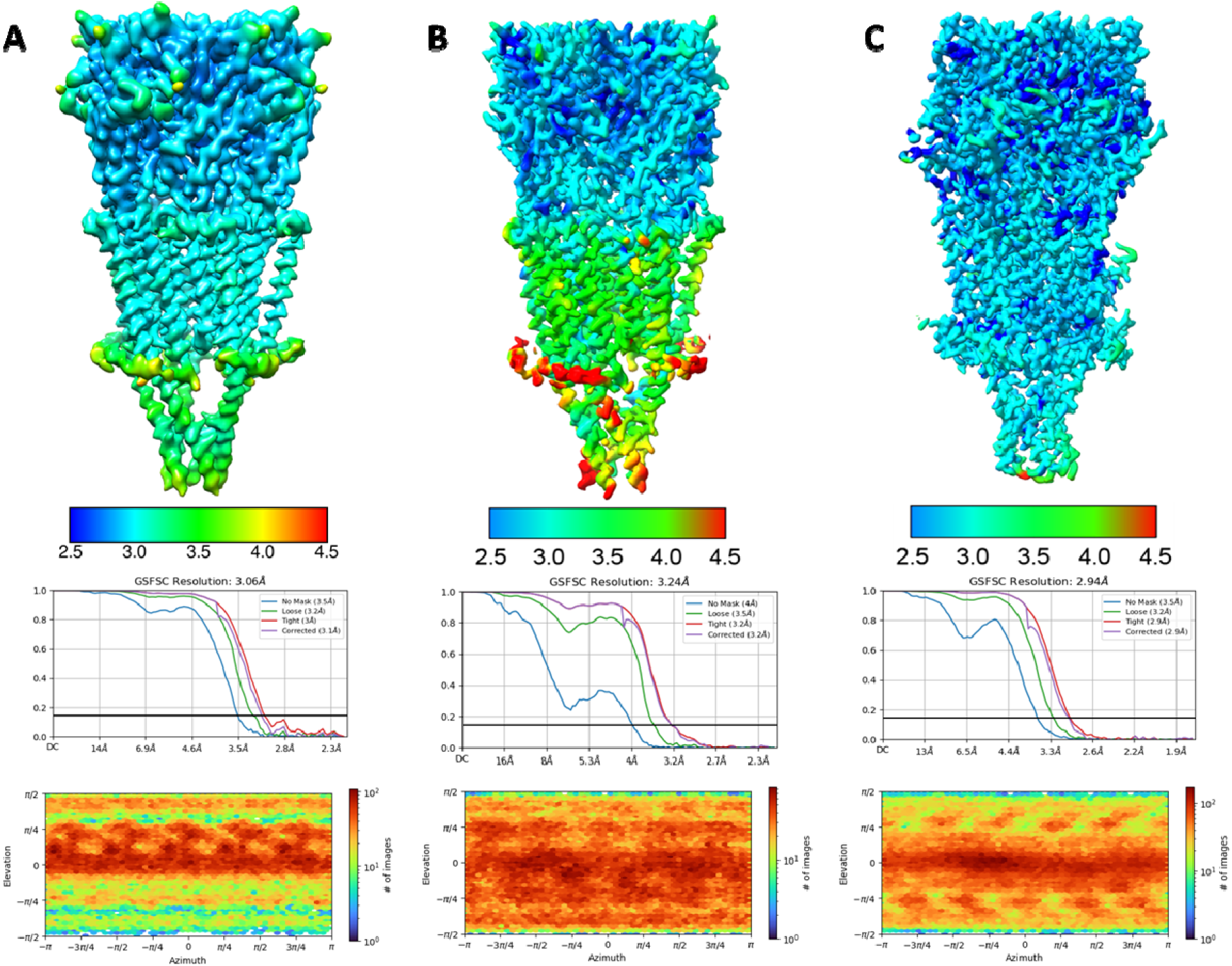
CryoEM processing results showing local resolution maps, GSFSC plots, and particle orientation distributions. **A)** ImI-α7-nAChR **B)** ImII-α7-nAChR **C)** ImII-mt-nAChR.

**Figure S6:**
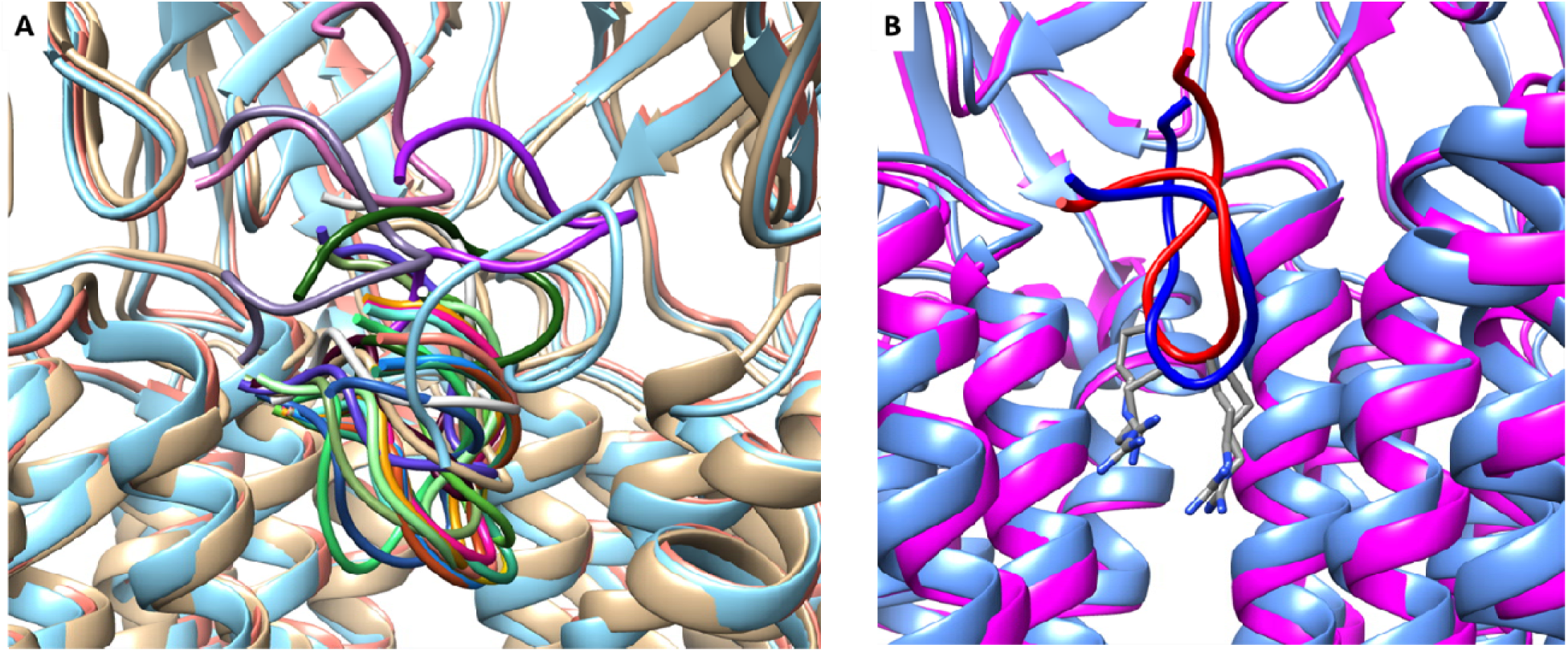
ToxDock results for the ImII-α7-nAChR complex and comparison to the cryoEM structure. **A)** Top 25 ToxDock predicted binding conformations overlaid. **B)** Overlay of the cryoEM ImII-α7-nAChR structure and the best ToxDock model. The CryoEM model is in cornflower blue and dark blue. The ToxDock model is magenta and red. The two pore-intruding arginine sidechains (Ctx-R6 and Ctx-R7) are shown.

**Figure S7:**
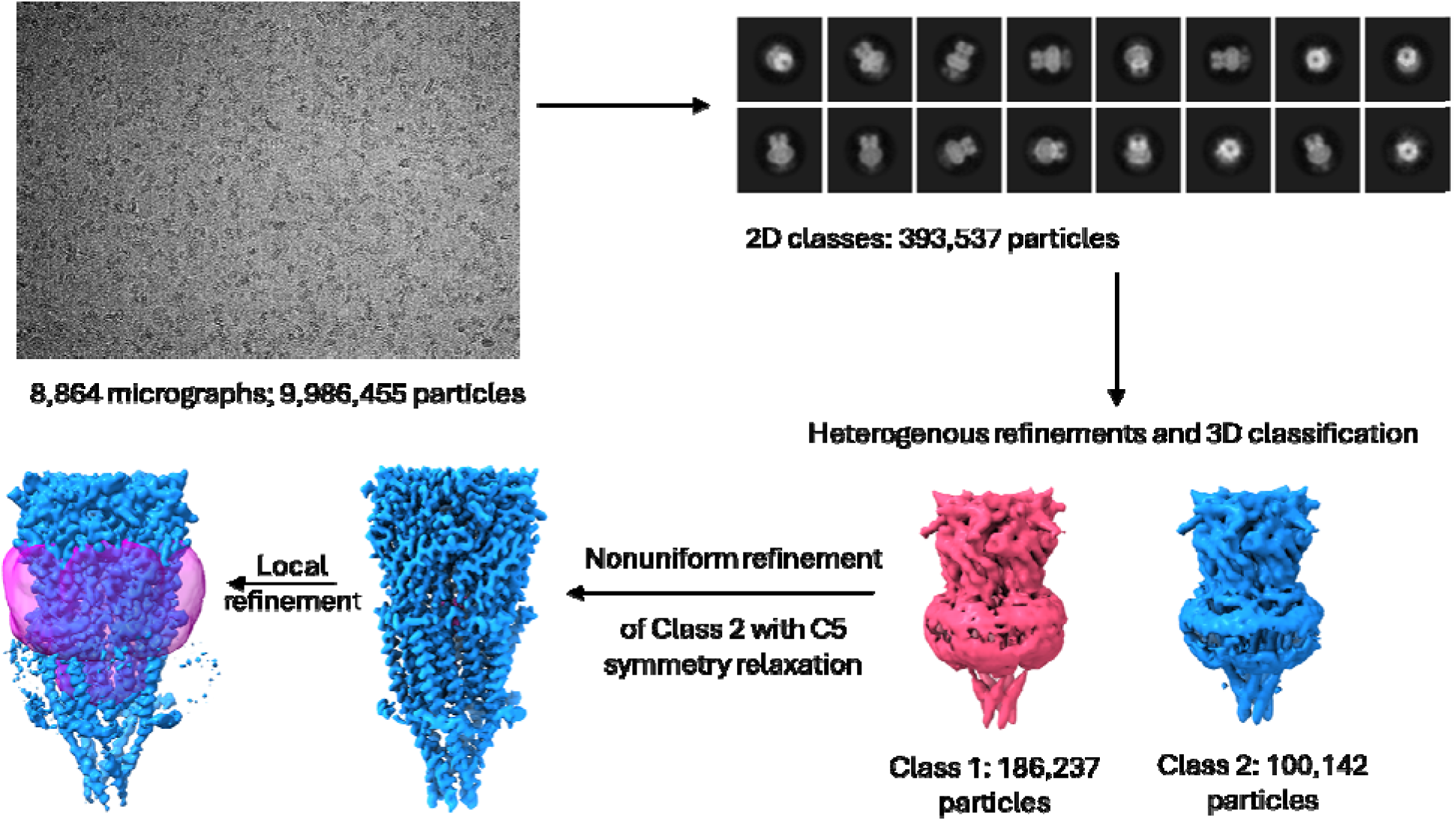
Data processing workflow with specific steps utilized for the ImII-α7-nAChR complex. After 2D classification and heterogenous refinement, 3D classification was performed with a focused and 2 major classes were identified, one with strong toxin density and one with weak toxin density. The class with clear toxin density was then subjected to non-uniform refinement with relaxed C5 symmetry. Subsequently a mask (purple toroid) was used for local refinement to improve the ImII density.

